# Reconsolidation-disruption diminishes spontaneous and stress-induced cocaine seeking

**DOI:** 10.1101/169839

**Authors:** Exton-McGuinness Marc TJ, Drame Mohamed L, Flavell Charlotte R, Lee Jonathan LC

## Abstract

**BACKGROUND:** There has been a recent surge of interest in exploiting the process of reconsolidation to weaken maladaptive memories, in the hope this will drive the next wave of innovation in psychotherapy. Reconsolidation normally functions to stabilise and maintain memories in the long-term, and is critical in enabling memory updating. However, this process can be disrupted pharmacologically to weaken memories, or harnessed to allow destructive interference of a memory trace. Work has already begun to exploit this mechanism to disrupt pavlovian fear memories in the treatment of maladaptive anxiety and threat processing, and additionally being able to target instrumental memories may provide further clinical benefit.

**METHODS:** Expanding our rat intravenous (i.v.) self-administration paradigm, we tested whether disruption of instrumental memory reconsolidation with the noncompetitive N-methyl-D-aspartate receptor (NMDAR) antagonist MK-801 could reduce relapse of cocaine seeking in response to stress, drug-priming or presentation of a drug-associated cue.

**RESULTS:** Spontaneous responding for i.v. cocaine was reduced by reconsolidation-disruption. Furthermore, responding was not rescued by pharmacologically-induced stress. However, responding was restored following systemic administration of the drug, or presentation of a drug-associated cue.

**CONCLUSIONS:** These data are consistent with hypothesis that there exist multiple ‘routes to relapse’, and suggest that at least some of these routes could be blocked by reconsolidation-disruption. This work provides important proof-of-principle that reconsolidation based therapies are a viable means of reducing the rates of relapse in substance use disorders.

## INTRODUCTION

Memories are constantly evolving through a constructive process that serves to update their content. One core mechanism of this is reconsolidation (1, 2). Following appropriate retrieval, a memory can be destabilized enabling the editing of its content. To persist in long-term memory, the updated trace must subsequently be restabilized, requiring activity at N-methyl-D-aspartate receptors (NMDARs), gene expression, and protein synthesis (3). Pharmacologically impairing the restabilisation of a memory during the reconsolidation phase can induce amnesia. This finding has generated much interest in treating disorders underpinned by maladaptive memories (4–6).

Substance use disorders (SUDs) are chronically relapsing, each bout of relapse often precipitated by exposure to stress or drug-associated cues. These ‘routes to relapse’ can be attributed to the underlying reward memories that support drug-seeking behaviours (5). It has been suggested that the formation of a maladaptive ‘habit’ memory following substance abuse (7) supports the underlying behavioural compulsion to seek drugs, regardless of consequences, that characterises the state of addiction (8). Importantly, both instrumental (operant) associations and pavlovian conditioned stimuli (CSs) contribute to the performance and maintenance of drug-seeking behaviours (9, 10). Weakening these reward memories by disrupting their reconsolidation may be a novel way to reduce rates of relapse.

It is now well-accepted that reconsolidation of pavlovian cue-drug memory can be disrupted using systemic NMDAR antagonism, thereby disrupting control of drug-seeking by pavlovian stimuli (11, 12). In human populations, weakening these pavlovian associations has successfully reduced drug cravings in abstinent addicts (13). Importantly for prospective treatment, these interventions do not impinge upon the underlying instrumental ‘habit’ memory which is believed to facilitate the compulsion to take drugs.

We recently demonstrated reconsolidation-disruption of appetitive instrumental memory with the NMDAR antagonist MK-801, using both sucrose and cocaine reinforcement (14). Here we extended our work on intravenous (i.v.) self-administration by prolonging the period of drug-taking, and investigated: 1) whether reconsolidation-disruption of instrumental cocaine memory can reduce drug-seeking after prolonged exposure; and 2) whether drug-seeking following reconsolidation-disruption could be reinstated by stress, re-exposure to the drug, or presentation of a drug-associated cue.

## METHODS AND MATERIALS

Subjects were 84 experimentally naïve male Hooded-Lister rats (Charles River, UK) weighing 250-350g (median 275g) at the beginning of the experiment. Rats were housed in quads in individually ventilated cages with aspen chip bedding on a 12hr light/dark cycle (lights on 0700). Environmental enrichment was available in the form of wood chew blocks and paper house. Food and water were provided *ad libitum.* Experimental sessions took place 0800-1200 each day. At the end of the experiment all animals were humanely killed via a rising concentration of CO_2_. All procedures were carried out in accordance with the UK Animals (Scientific Procedures) Act 1986, Amendment Regulations 2012 (PPL P8B15DC34).

### Surgical procedures

Drinking water was supplemented with the broad-spectrum antibiotic Baytril for 7d, beginning 3d pre-operatively. Rats were anaesthetised using isoflurane (5% induction, 2-3% maintenance), and administered peri- and post-operative buprenorphine; their diet was also supplemented with the non-steroidal anti-inflammatory Carprofen for 1d pre-operatively, and 3d post. Rats were allowed a minimum of 7d recovery, during which they were singly housed on Puracel bedding; rats were rehoused in quads at the start of experimental procedures.

Surgeries were carried out aseptically (15) during which rats were implanted with chronic indwelling jugular vein catheters (Polyurethane Intravascular Tubing; Instech, PA) aimed at the left vena cava, secured with silk suture, and exteriorised on the dorsum with a small plastic implant (PlasticsOne, VA; 313-000BM-15-5UP/1/SPC) secured to the skin with a 1 inch mesh.

### Drugs

Cocaine (Macfarlane Smith Ltd, UK) was dissolved in sterile saline to a concentration of 2.5 mg/ml; i.v. infusions of 0.1ml over 5.6 seconds could be obtained during training and reactivation. Infusion dosage was based upon established literature (16). For cocaine-primed reinstatement, 10 mg/kg of cocaine solution was injected intraperitoneally (i.p.) immediately before the behavioural session; this dose is known to reinstate lever pressing for cocaine (17). MK-801 (AbCam, UK) was dissolved in sterile saline to a concentration of 0.1 mg/ml. 30 minutes prior to memory reactivation rats were injected i.p. with 0.1mg/kg of MK-801 or saline vehicle. This dose is established to disrupt instrumental memory (14, 18). Injections were assigned systematically by cage, randomly within each cage. Yohimbine (Sigma-Aldrich, UK) was dissolved in sterile saline to a concentration of 1.25 mg/ml. For stress-induced reinstatement, yohimbine was administered 30 minutes prior to testing (1.25 mg/kg). This dose is established to reinstate drug-seeking in cocaine settings (19, 20).

### Behavioural procedures

Behavioural sessions took place in 8 operant boxes (MedAssociates, VT), as described previously (14). Prior to each session catheters were flushed with heparinised saline (0.1 ml, 30 IU/ml). Catheters were then connected to an infusion pump (MedAssociates, VT) and secured with a spring tether.

#### Training

Rats were trained to self-administer cocaine for 10d, on a fixed-ratio schedule. Two levers were extended into the chamber, one assigned the ‘active’ reinforced lever. Lever assignments were made systematically prior to the start of training. Active responses triggered delivery of a single cocaine infusion and a 20-second illumination of a light conditioned stimulus (CS) above the active lever, during which the houselight went out. Both levers remained extended throughout the session, and inactive responses had no consequence. A 20-second timeout was enforced between infusions. Training sessions lasted 90 minutes, or until 30 infusions.

#### Reactivation

48 hours after the final training day, rats were injected i.p. with MK-801, or saline, 30 minutes prior to a variable-ratio (VR5) reactivation session. VR5 required a random number of active lever presses to gain an infusion (mean: 5, range: 1-9). Reactivation lasted 20 minutes, or until 20 infusions.

#### Testing

The following day, responding was tested in a 90-minute extinction session. Levers were extended throughout, however neither the drug nor drug-paired CS was presented. Separate cohorts of rats were tested for reinstatement with yohimbine-induced stress, re-exposure to the drug, or the drug-associated cue (1 sec contingent CS).

### Statistical analysis

Data are represented as mean ± SEM, and analysed by ANOVA using JASP (21). Results p<0.05 were deemed significant. For acquisition data, Training day was included as a factor; where appropriate a Greenhouse-Geisser correction was applied. Lever responses (active/inactive) were included as a factor for each analysis, with assignment (left/right) included as a cofactor. Drug treatment was assigned pseudo-randomly to produce two groups similarly performing during training; total infusions during training was included as a covariate for analysis of reactivation and test data. Post-hoc comparison was performed on saline control groups to assess any enhancement of responding by the different reinstatement conditions using one-way ANOVA. 6 rats were excluded due to problems with the catheters or equipment. 5 rats were excluded as statistical outliers (2 s.d. from mean). 2 rats were excluded due to a failure to learn the task (<25 total infusions).

## RESULTS

### Experiment 1: Disruption of reconsolidation

Rats learned to self-administer cocaine over 10d (Lever x Training: F_(4.8,106.0)_=13.544, p<0.001, η^2^=0.259), with no significant differences in responding between treatment groups (all F<1). At reactivation there was an acute overall reduction in responding following administration of MK-801 (Treatment: F(_1,21)_=7.499, p=0.012, η^2^=0.179; Lever x Treatment: F_(1,21)_=2.439, p=0.133, η^2^=0.073). When tested 24 hours later (Figure 1), rats previously treated with MK-801 made significantly fewer active lever responses (Lever x Treatment: F_(1,21)_=5.094, p=0.035, η^2^=0.163; Active lever [Saline vs. MK-801]: F_(1,21)_=6.019, p=0.023, η^2^=0.189; Inactive lever [Saline vs. MK-801]: F<1). Disruption of underlying lever pressing suggests instrumental memory was impaired by our manipulation.

**Figure 1.**
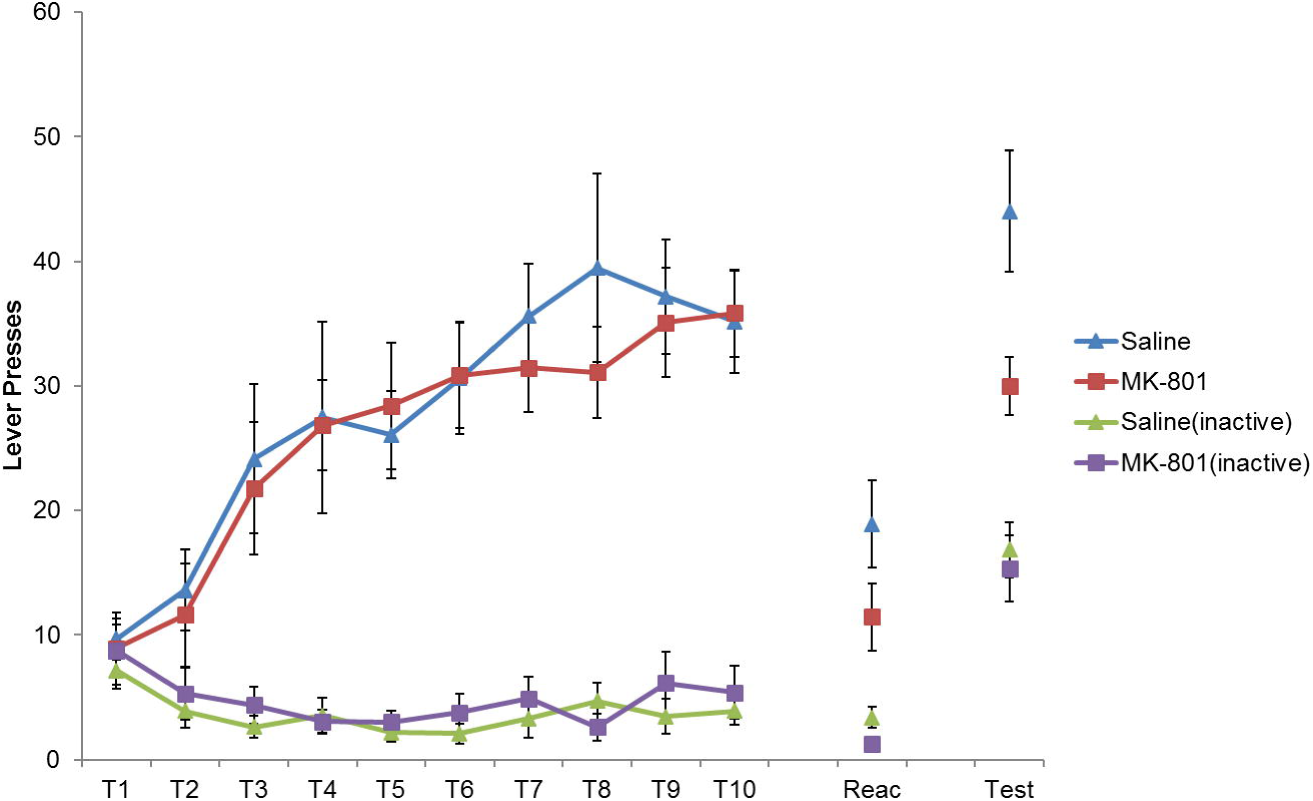
Long-term lever pressing was significantly impaired at test following administration MK-801 in conjunction with a reactivation session. Rats learned to self-administer cocaine over training days 1-10 (T1-T10). At reactivation, lever pressing was reduced by MK-801 administration. When tested 24 hours later, previously MK-801 treated rats remained significantly impaired in their responding. (Saline, n=13; MK-801, n=13).

### Experiment 2: Stress induced reinstatement

A separate cohort of rats was successfully trained as before (Lever x Training: F_(4.5,50.0)_=5.294, p<0.001, η^2^=0.281) with no significant group differences (Lever x Training x Drug: F(45,500)=1.063, p=0.389, η^2^=0.056; all other F<1). At reactivation, there was again an overall reduction in responding following injection of MK-801 (Treatment: F_(1,10)_=8.166, p=0.017, η^2^=0.333; Lever x Treatment: F_(1,10)_=4.622, p=0.057, η^2^=0.194). The next day, rats were administered i.p. yohimbine prior to testing (Figure 2). Responding was not recovered, and active responses remained significantly reduced in MK-801-treated rats (Lever x Treatment: F_(1,10)_=7.581, p=0.020, η^2^=0.360; Active lever [Saline vs. MK-801]: F_(1,10)_=15.617, p=0.003, η^2^=0.542; Inactive lever [Saline vs. MK-801]: F_(1,10)_=2.990, p=0.114, η^2^=0.224). This implies reconsolidation-disruption conferred resistance to stress-induced relapse.

**Figure 2.**
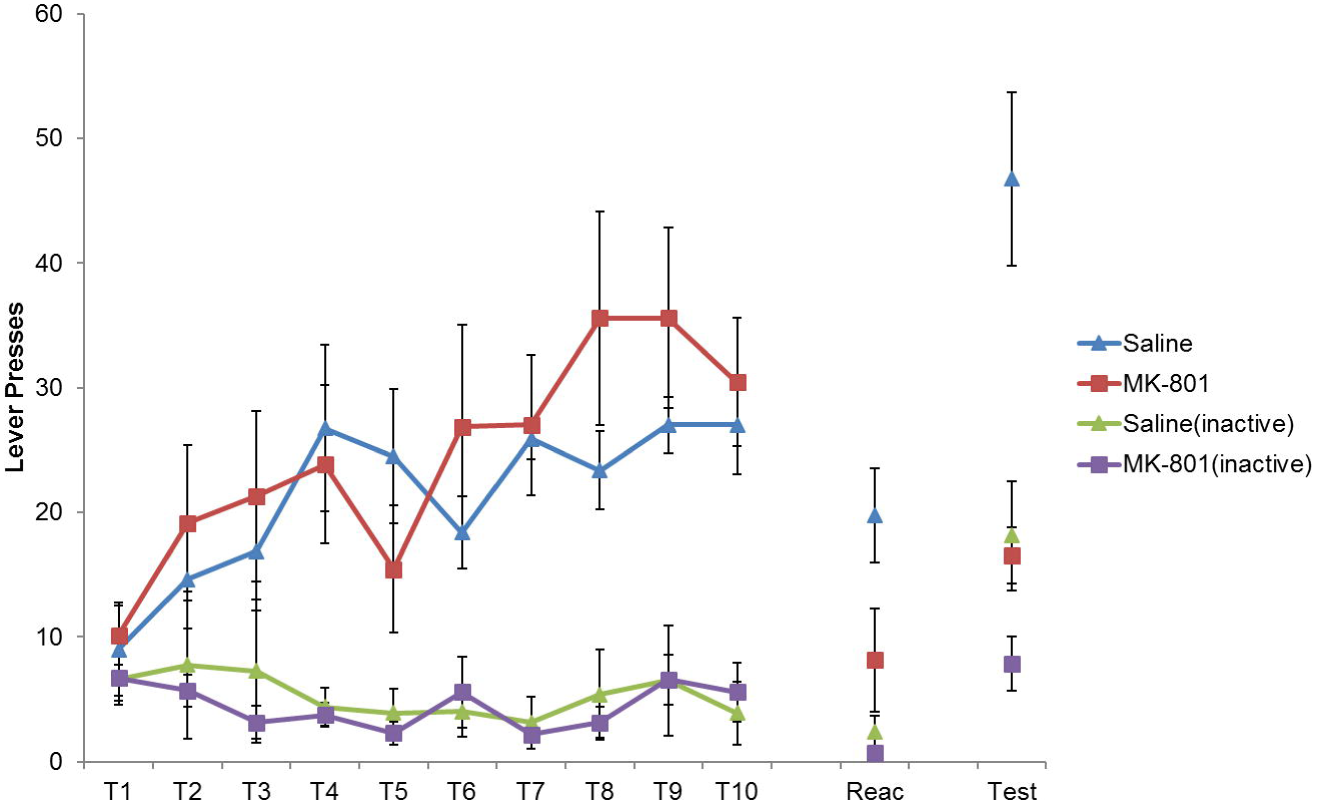
Induction of stress using the α-2 antagonist yohimbine failed to reinstate lever pressing following reconsolidation-disruption with MK-801. Rats learned to lever press for cocaine over training days 1-10 (T1-T10). At reactivation, lever pressing was acutely reduced by MK-801. Prior to testing rats were injected with yohimbine, however this did not recover performance and MK-801-treated rats remained impaired (Saline, n=8; MK-801, n=7).

### Experiment 3: Drug induced reinstatement

The cocaine-lever contingency was learned as before over 10d (Lever x Training: F_(4.1,5.28)_=3.628, p=0.011, η^2^=0.184) with no significant group differences (Lever x Treatment: F_(1,13)_=2.097, p=0.171, η^2^=0.022; Lever x Training x Treatment: F_(4.1,52.8)_=1.508, p=0.213, η^2^=0.077; all other F<1). At reactivation there was again a mild acute effect of MK-801 treatment (Treatment: F_(1,12)_=4.892, p=0.047, η^2^=0.248; Lever x Treatment: F_(1,12)_=3.835, p=0.074, η^2^=0.184). 24 hours later, rats were given a priming injection of cocaine immediately prior to testing (Figure 3). Cocaine-priming appeared to rescue the previous MK-801-deficit, with no significant differences in responding between treatment groups (all F<1).

**Figure 3.**
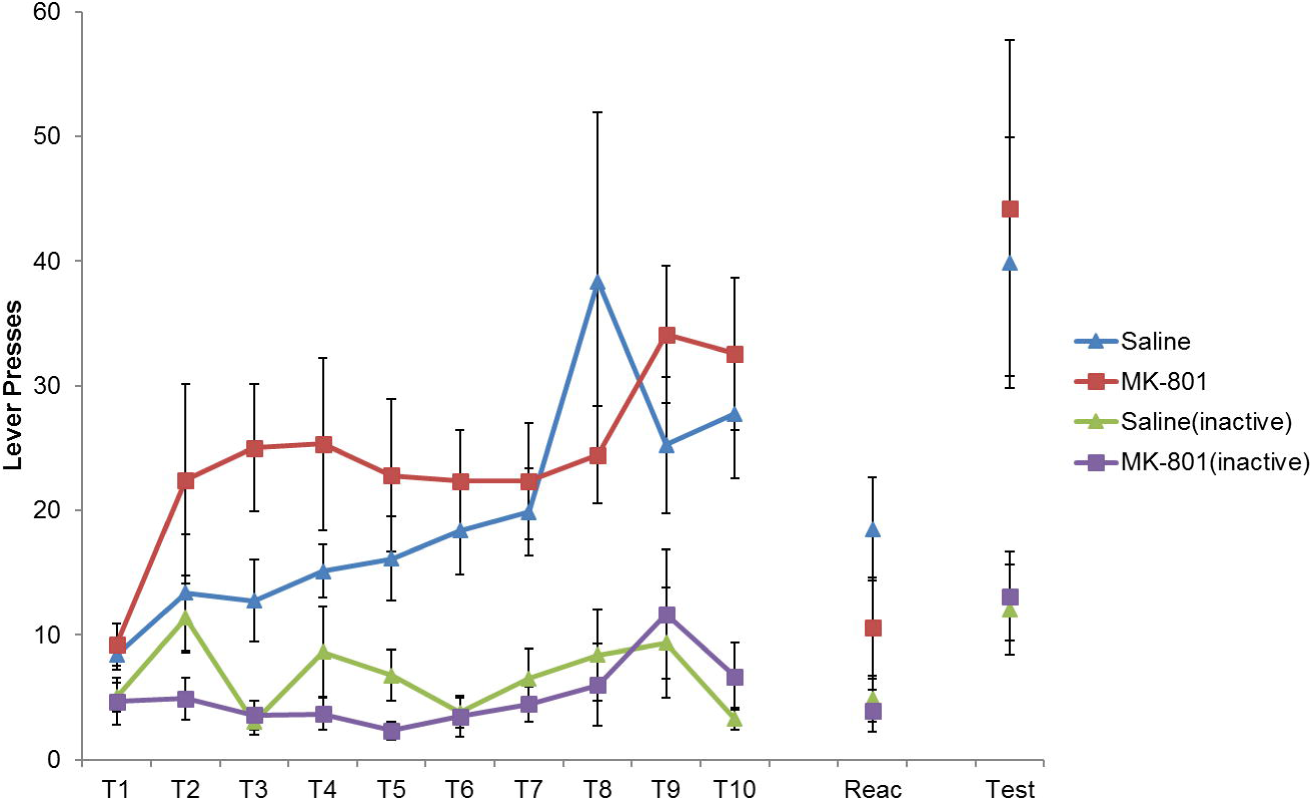
Administration of a cocaine priming injection successfully rescued lever pressing on the active drug-lever, following MK-801 induced disruption of reconsolidation. Rats acquired cocaine seeking successfully over training days 1-10 (T1-T10). At reactivation, lever pressing was slightly reduced by MK-801 administration. Rats were injected with cocaine immediately prior to testing, which recovered the MK-801-impaired rats to control levels of performance (Saline, n=8; MK-801, n=9).

### Experiment 4: CS induced reinstatement

One final cohort was trained to lever press for cocaine; again rats learned to discriminate between levers (Lever x Training: F_(2.6,23.8)_=5.161, p=0.009, η^2^=0.316), with no significant group differences (all F<1). During reactivation responding was mildly, although not significantly, reduced in MK-801-treated rats (Treatment: F_(1,8)_=3.047, p=0.119, η^2^=0.164; Lever x Treatment: F_(1,8)_=1.373, p=0.275, η^2^=0.061). Long-term persistence of responding was tested the next day, during which active responses triggered a brief CS presentation (Figure 4). CS delivery was sufficient to restore responding of MK-801-treated rats to control levels (all F<1). This indicates both that CS-drug memory was spared by our intervention, and that this memory can recover drug-seeking behaviour following instrumental reconsolidation-disruption.

**Figure 4.**
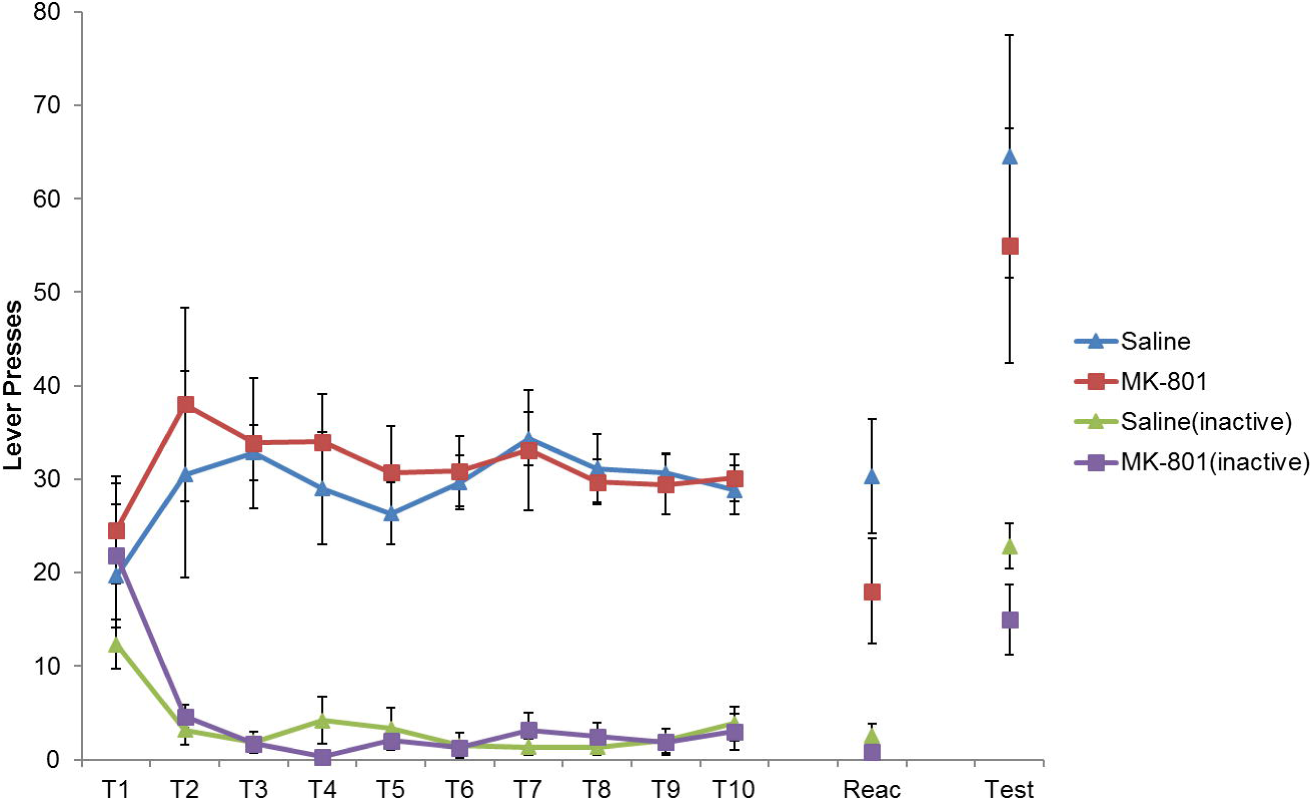
Presentation of a drug-paired CS maintained lever pressing, following prior disruption of reconsolidation with MK-801. Rats acquired cocaine seeking successfully over training days 1-10 (T1-T10). At reactivation, lever pressing was slightly reduced by MK-801, although this was not significant. At test, active lever responses triggered a 1-second CS presentation; this recovered lever responding on the active lever to control levels. (Saline, n=6; MK-801, n=7).

## DISCUSSION

Our data demonstrate spontaneous drug-seeking behaviour can be attenuated by reconsolidation-disruption of operant memory. In line with our past findings, systemic NMDA receptor antagonism in conjunction with a change in instrumental contingency directly weakened later lever pressing behaviour. However, pavlovian cue-drug memory was spared. Interestingly, responding was reinstated by a cocaine priming injection but not by yohimbine-induced stress, suggesting these two forms of reinstatement act via distinct mechanisms.

We have previously shown changes in reward contingency destabilize instrumental sucrose memory, successfully disrupting reconsolidation with MK-801 (18); similar findings were observed for weakly-trained cocaine memory (14). Here we extended the length of cocaine self-administration, and successfully destabilized the underlying drug memory by implementing an adapted protocol in which our VR5 reactivation was given 48 hrs after training; delaying the reactivation session appears to facilitate destabilisation in sucrose settings (*unpublished observations*). Given our previous demonstration that the decrease in lever pressing is due to disruption of instrumental memory (18), it is highly likely the present reduction in cocaine seeking is similarly caused by an impairment in instrumental memory.

While MK-801 did have a mild acute effect during reactivation, it is unlikely this caused any long-term inhibition of lever pressing: recovery of responding with drug or cue exposure was complete and there was no evidence of motor impairment. The acute effect of MK-801 on performance may be related to its ability to itself reinstate cocaine seeking (22); however, this cannot explain our data as long-term performance was impaired, not recovered. Finally, MK-801 does not appear to have affected the long-term incentive value of cocaine as responding was recovered to control levels under certain conditions.

Memory disruption was specific to spontaneous lever pressing; pavlovian cue-drug memory was spared as evidenced by the ability of response-contingent CS presentation to rescue performance. This highlights the specificity of reconsolidation manipulations to the active memory trace. Operationally it appears which memory is destabilized depends upon what functional changes are made at reactivation (or what new information is presented): changing the response-reward contingency destabilizes instrumental memory, while altering cue-reward contingency causes pavlovian memories to undergo reconsolidation (23). It is notable that, in contrast to our data here, pavlovian reconsolidation disruption does not impinge upon the underlying baseline lever pressing (11, 23). This raises an important issue for translation to humans, as destabilisation of pavlovian and instrumental memories may be mutually exclusive. Further study is needed to determine whether these memories can be disrupted in tandem.

A key question arising from our data is how the CS recovered cocaine-seeking, given instrumental memory appeared to be impaired. It is well-established that CSs are able to act as conditioned reinforcers (24) and induce states of drug craving (25, 26). However, it is not immediately clear how this would recover lever pressing following disruption of instrumental memory. It may be that the CS allowed reacquisition via its capacity as a conditioned reinforcer, although response rates observed following this type of CS-mediated learning are typically very low (27, 28) and thus likely insufficient alone to account for the near total rescue of behaviour by the CS.

Another possibility is that the CS induced a drug craving which drove responding; however, this would require instrumental memory to be at least partially intact. It may be the reconsolidation-disruption significantly weakened the instrumental drug-seeking memory, but did not erase it completely. There is some evidence for this (as performance on the active lever was generally higher than the inactive), however the assertion memory was sufficiently preserved to allow reinstatement via central emotional states and cravings is incompatible with the lack of reinstatement with yohimbine (see below). Moreover it is unclear why recovery of responding should be to control levels.

A final hypothesis is that the CS supported lever pressing independently of instrumental memory. It has been observed that high levels of lever pressing can be supported by drug-paired CSs (29). One interpretation of these findings is that motivational processes become associatively coupled to operationalised habits through pairing a conditioned reinforcer with the response; termed an ‘incentive habit’ (10). If such learning took place then it may explain how the CS was able to recover responding to similar levels as controls. It is worth emphasising that ‘incentive habits’ bear much similarity to pavlovian conditioned responses (David Belin, *personal communication*), highlighting that this behaviour is distinct from instrumental responding. This has important implications for our understanding of SUDs, as maladaptive incentive processes likely play a key role in the progression of addiction.

Notably, reconsolidation-disruption of lever pressing for cocaine was not reinstated by induction of pharmacological stress. Yohimbine has been observed to recover lever pressing from extinction, for cocaine (20), nicotine (30) and heroin (31) at the doses used here. These data would imply that cocaine-seeking could no longer be motivated by changes in central motivational and/or emotional state, consistent with reconsolidation-disruption of instrumental memory. However it is worth noting that yohimbine, and stress more generally, appears to have strong effects upon CS-mediated forms of reinstatement (20, 31, 32). Thus, while stress in isolation does not appear to be sufficient to recover the impaired instrumental responding, it may still act synergistically to enhance cue-induced relapse.

In contrast to yohimbine, a cocaine priming injection did reinstate lever pressing behaviour in a similar fashion to re-exposure to the CS. Given the lack of effect with yohimbine, the underlying mechanisms driving these two forms of reinstatement are likely distinct (33), or at least cocaine priming has additional effects to precipitate relapse. It seems unlikely the effect of cocaine priming was mediated via stress, or cue-reactivity (no discrete cues were presented at test).

One interpretation is cocaine-priming induced craving for the drug (26, 34), similarly to the CS. Indeed, an interesting question that arises is whether the mechanisms supporting cue- and drug-induced reinstatement are similar in our paradigm. During training both the CS and the drug itself were temporally coincident, and would have both been predictive of a drug-seeking state (8, 35); thus both could have become conditioned to act as pavlovian CSs to trigger a drug-specific cocaine craving, *vis a vis* two process theory (36). Similar counter-arguments apply as before. If relapse occurred via a craving mechanism, then instrumental memory must have been at least partially intact, but this is inconsistent with our initial findings and the lack of a yohimbine-induced reinstatement effect. However, if cocaine-presentation itself truly became a CS then it may have supported an ‘incentive habit’ as previously suggested for our cue-induced recovery. Moreover, if cocaine-priming acted as a CS this may explain the differences between this effect and the lack of effect with yohimbine. Again, this raises the possibility that stress and drug re-exposure might interact during relapse (see above).

An alternative interpretation of our drug-primed reinstatement effect is that the instrumental memory was fully recovered, perhaps via a state dependent retrieval mechanism (37, 38). From this viewpoint, the cocaine would have acted as a retrieval cue to prime lever pressing. Although this account could explain differences observed between yohimbine and cocaine-induced reinstatement, it is not entirely satisfactory as if retrieval were more generally dependent upon cocaine presentation then performance of saline-treated rats should have been enhanced by cocaine-priming; post-hoc comparisons of active responses in saline control groups provided no significant evidence to support this assertion (F<1, p=0.492, η^2^=0.067). Reconsolidation-disruption of the underlying instrumental memory remains the most parsimonious conclusion which accounts for our findings in all four experimental conditions.

Our data provide strong evidence that the instrumental memories which support lever pressing for cocaine can be disrupted to reduce drug seeking behaviour. Responding was resistant to reinstatement via pharmacological stress, however pavlovian mechanisms which were spared by our manipulation could still recover responding following presentation of drug-associated CSs. Importantly, it has long been established that the reconsolidation of pavlovian drug-cue memory can be disrupted (11, 12, 23), as can the motivational memories that support conditioned withdrawal (39). Combined with our data presented here, reconsolidation-disruption has been demonstrated for many different components that support drug-seeking and relapse. Together these works lay the groundwork for the development of future reconsolidation based treatments for SUDs.

Our data provide important proof-of-principle that reconsolidation-disruption is a viable mechanism for reducing spontaneous drug seeking. Our disruption appeared to impair the underlying operant response for drug, which was not reinstated by yohimbine-induced stress. However, responding did recover following re-exposure to the drug-itself or a drug-paired cue. Reconsolidation-disruption of reward memories offers a powerful tool to meaningfully reduce drug seeking in SUDs, however erasing both the pavlovian and instrumental memories which support addiction is necessary to provide the most effective anti-relapse intervention.

## ACKNOWLEDGEMENTS

The authors wish to acknowledge the assistance of David Barber with data collection.

